# Hunter-gatherer oral microbiomes are shaped by contact network structure

**DOI:** 10.1101/2022.05.03.489993

**Authors:** Federico Musciotto, Begoña Dobon, Michael Greenacre, Alex Mira, Nikhil Chaudhary, Gul Deniz Salali, Pascale Gerbault, Rodolph Schlaepfer, Leonora H. Astete, Marilyn Ngales, Jesus Gomez-Gardenes, Vito Latora, Federico Battiston, Jaume Bertranpetit, Lucio Vinicius, Andrea Bamberg Migliano

**Author notes:** equal contribution (Jaume Bertranpetit, Lucio Vinicius, Andrea Bamberg Migliano). equal contribution (Federico Musciotto, Begoña Dobon).

## Abstract

Ancestral humans evolved a complex social structure still observed in extant hunter-gatherers. Here we investigate the effects of extensive sociality and mobility on the oral microbiome of 138 Agta hunter-gatherers from the Philippines. Comparisons of microbiome composition showed that the Agta are more similar to Central African Bayaka hunter-gatherers than to neighboring farmers. We also defined the Agta social microbiome as a set of 137 oral bacteria (only 7% of 1980 amplicon sequence variants) significantly influenced by social contact (quantified through wireless sensors of short-range interactions). We show that interaction networks covering large areas, and their strong links between close kin, spouses, and even unrelated friends, can significantly predict bacterial transmission networks across Agta camps. Finally, more central individuals to social networks are also bacterial supersharers. We conclude that hunter-gatherer social microbiomes, which are predominantly pathogenic, were shaped by evolutionary tradeoffs between extensive sociality and disease spread.

## Introduction

Hominins have significantly diverged from other African apes regarding social behaviour and structure^1^. Compared to polygynous mating and male philopatric residence patterns typically found in chimpanzees, bonobos and gorillas, archaeological and ethnographic evidence point to a stepwise emergence of features such as pair bonding, multilocal residence, high mobility between residential camps and increased co-residence with unrelated individuals^2,3^. Such traits were the foundations of multilevel social structuring appearing in ancestral *Homo sapiens* and possibly earlier hominins. The niche of extant hunter-gatherers may offer a window into past human adaptations as it still exhibits features prevalent before the advent of agriculture, such as a high-quality diet including meat and tubers, and multilevel sociality. Multilevel organization results in interconnected social networks covering large areas and multiple residential camps^4^, and in frequent interactions between individuals differing by sex, age and relatedness level. Interconnected networks may have accelerated the evolution of cultural innovations in humans compared to other apes^5,6^. However, efficient networks may also facilitate the spread of infectious diseases^7^, potentially affecting the structure and composition of hunter-gatherer microbiomes. Previous studies have investigated the role of diet, ecology and environment in hunter-gatherer oral, gut and milk microbiomes^8–16^ and revealed higher oral microbiome diversity in hunter-gatherers than in farming populations^17^. However, they have not been able to isolate the contribution of high sociality and individual mobility to microbial transmission from other factors such as shared environments or diet. Although the more fluid and complex sociality of hunter-gatherers results in high levels of camp coresidence^2^, cooperation and social interactions among unrelated individuals^18^, its potential effects on microbiome transmission have been mostly neglected. We conducted a comprehensive investigation of the oral microbiome of Agta hunter-gatherers to analyse the specific effect of sociality and social network structure on the composition of the Agta oral microbiome, with a companion article examining the separate role of environmental (diet) and biological (age, sex, host genotype) factors^19^.

We obtained both oral microbiome sequences and high-resolution social network data from the same 138 Agta hunter-gatherers from the Philippines. We also collected oral microbiome data for 21 Bayaka hunter-gatherers from the Congo, and 14 Palanan farmers neighboring the Agta territory. We sequenced the 16S rRNA region and identified 6409 amplicon sequence variants (ASVs)^20^, later reduced to 1980 ASVs (with at least 10 counts and present in at least two individuals), to detect fine-scale variation between individuals. We also collected data on proximity interactions and social networks using radio sensor technology recording close-range dyadic interactions every two minutes for 5-7 days^6,18^ from four Agta camps, and from two longer multi-camp experiments (interactions recorded every hour for one month). Proximity data were supplemented with information on household composition, kinship and affinal relationships from all Agta individuals.

Our extensive dataset on oral microbiome composition and social interactions from the same individuals allowed us to investigate in more depth the possible effects of sociality on oral microbiome transmission and composition in Agta hunter-gatherers. Our aims were to investigate the roles of hunter-gatherer niche and geography on oral microbiome diversity in hunter-gatherers from two continents and a neighboring farming population from the Philippines; to determine which fraction of the Agta oral microbiome specifically responds to levels of social interaction; to identify levels of pathogenicity of the oral microbiome transmitted through social contact; to investigate any potential tradeoffs between increased sociality and the spread of infectious disease; and to verify potential tradeoffs at individual level by testing whether ‘hyper social’ individuals also shared more bacteria. In the following, we provide evidence that the oral microbiome of extant hunter-gatherers was partially shaped by tradeoffs between extensive sociality and the spread of infectious disease.

## Results

### Hunter gatherer niches shape the oral microbiome

To investigate the contributions of lifestyle versus environment to the hunter-gatherer oral microbiome, we compared the Agta (n=138) to smaller samples of Bayaka hunter-gatherers from the Congo (n=21) and neighboring Palanan farmers from the Philippines (n=14) (see Methods). Both Agta (mean of 252.1±90 ASVs per individual) and Bayaka (280.1±83) exhibited significantly more ASVs than Palanan farmers (163.4±34) (P<0.0001; Figure 1a), and higher levels of ASV diversity as measured by Faith’s Phylogenetic Diversity index (Figure 1b). Comparisons based on the total set of ASVs in each population (controlling for differences in sample size through subsampling) revealed that the Agta shared more bacteria with African Bayaka (471.2±33.9) than with neighboring Palanan farmers (423.4±23.8) (Figure 1c). Finally, Agta and Bayaka resampled groups showed respectively 651.5±93.7 and 688.1±61.9 exclusive ASVs, against only 285.1±16.9 in Palanan farmers (Figure 1d). In summary, the two hunter-gatherer populations show higher microbiome diversity and uniqueness than Palanan farmers, consistent with findings that farming significantly reduced gut microbiome diversity^21^. Results therefore demonstrate the precedence of niche over geography in shaping hunter-gatherer oral microbiomes.

**Figure 1.**
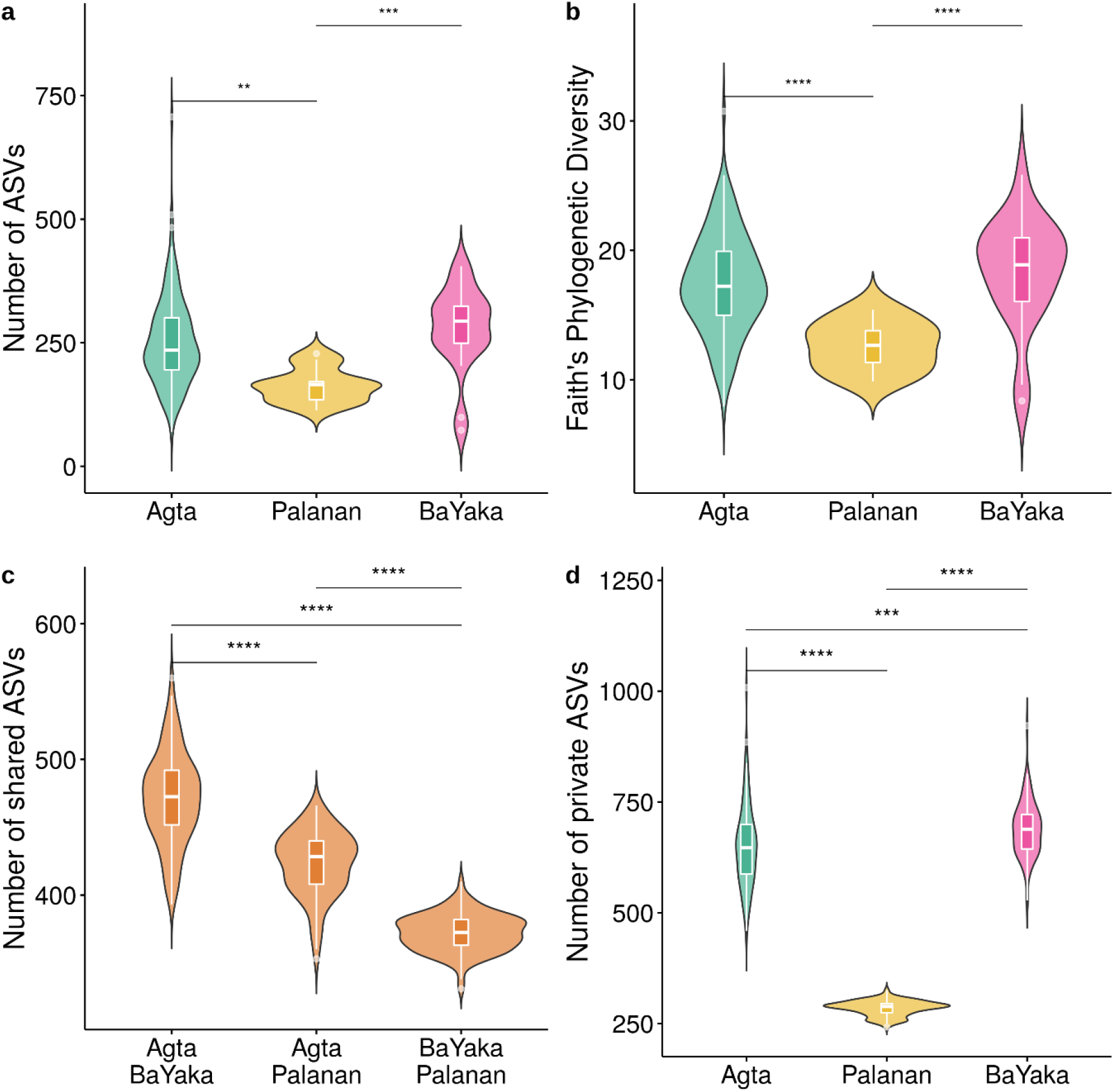
Oral microbiome diversity in Agta hunter-gatherers, neighbouring Palanan farmers, and Bayaka hunter-gatherers. a) Number of ASVs in the Agta (n=138), Bayaka (n=21) and Palanan (n=14). b) Oral microbiome diversity assessed by Faith’s Phylogenetic Diversity index accounting for ASV phylogenetic distances (Agta=17.52±3.83; Bayaka=18.45±4.10; Palanan farmers=12.62±1.86); c) Shared ASVs between populations, estimated by randomly sampling 10 individuals from each population (averaged over 100 permutations); d) Exclusive ASVs per individual, estimated by randomly sampling 10 individuals from each population (100 permutations). Boxplot midlines represent medians, and box limits represent first and third quartiles (****: FDR-adjusted P<0.0001; ***: P<0.001; **: P<0.01).

### The social microbiome is a socially transmitted fraction of the oral microbiome

Primate social ‘pan-microbiomes’ were recently defined as the totality of microorganisms present in a host population or species^22^, but this definition also includes microorganisms acquired due to common diet or environment. Here, we define the ‘social microbiome’ as the oral microbiome specifically transmitted through social interactions. To identify the socially transmitted fraction of the Agta oral microbiome, we used the contact network recorded through radio sensor devices and split all Agta dyadic social interactions into a strong (top 25% from the distribution of dyadic link weights) and a weak set (the remaining 75% links; see Methods). We then tested for differences in the proportion of each of the 1980 ASVs between the strong and weak sets. We identified 137 ASVs (7% of the Agta oral microbiome; see Supplementary Figure 1 and Supplementary Table 1) whose presence was significantly higher in the strong set, and therefore statistically associated with higher frequencies of social interactions. In the following we investigate the transmission patterns and composition of the hunter-gatherer social microbiome.

### The hunter-gatherer social microbiome is predominantly pathogenic

Human sociality is associated with multiple fitness benefits, including increased reproductive success^7^. reputation^23^. food sharing^24^, cooperation^3,25^ and cultural transmission^26^, but may also facilitate pathogen transmission^7,18^. In our dataset, from the 18 ASVs that could be classified at species level, 14 are socially transmitted, 9 of which (64.3%) are typically pathogenic, and 10 (71.4%) are typically oral. By contrast, all four non-socially transmitted species were non-pathogenic and typically oral.

We were able to classify 1886 of the 1980 ASVs at genus level, resulting in 36 socially (those representing ASVs included in the social microbiome) and 62 non-socially transmitted genera (the remaining ones). Among the social genera, 61.8%were classified as typically or exclusively pathogenic (21 out of 34; two genera could not be classified), against only 16.4% among non-socially transmitted genera (10 out of 61; one genus could not be classified). We identified many socially transmitted genera either typically (*Aggregatibacter*, *Capnocytophaga*) or uniquely (*Corynebacterium*) associated with dental plaque formation, gingivitis and calculus, the full red complex of periodontal disease (*Porphyromonas, Treponema* and *Tannerella*), and other potential periodontal pathogens (*Prevotella, Desulfobulbus, Fusobacterium*)^27–30^. The classification of bacterial genera as pathogenic is not unequivocal for those cases where some species within the genus can be pathogenic and others non-pathogenic. Thus, practical criteria were applied in these cases for assigning a genus to the pathogenic group (see Methods). Following these criteria, the social microbiome clearly has a higher proportion of pathogenic organisms than the non-socially transmitted portion.

We also found pathogenic bacteria typical of the gut (*Rickenellaceae*), non-human environments (*Tetragenococcus*, *Comamonas*), respiratory tract (*Staphylococcus*, *Moraxella*, *Streptococcus pneumoniae*), and both urogenital and respiratory tracts (*Mycoplasma*), suggesting that their spread may be facilitated by oral transmission^31–33^. In summary, the predominantly pathogenic nature of the social microbiome suggests a trade-off between benefits of hunter-gatherer sociality and costs associated with disease transmission.

### Hunter-gatherer multilevel social structure shapes social microbiome sharing

Hunter-gatherer sociality is characterised by specific interaction channels not found in non-human apes, such as long-term pair bonding and households, extended families, friendships among unrelated individuals, and frequent between-camp relocation. We estimated the effect of relatedness level, residence camp and friendships on the probability of sharing socially transmitted bacteria. First, we built a bacterial sharing network, where the weight of each Agta dyadic link is given by how many of the 137 social bacteria are shared by the two Agta individuals (rather than by the strength of its social bond, as in the social network). Next, we classified all dyadic links in this network into: i) levels of kinship (mother-offspring, father-offspring, siblings: *r*=0.5, other kin: *r*=0.25 or r=0.125, non-kin: *r*=0.0625 or lower, spouses, friends: defined as non-kin at the top 25% distribution of social dyadic weights, and other non-kin) and ii) residence (same or different camp, same or different household). Finally, we compared the mean weight of each type of dyadic link in our bacterial sharing network to its mean weight in a sample of 1000 networks of the same size and topology, but where the dyadic classification was randomised (Figure 2 and Supplementary Tables 2 and 3).

**Figure 2.**
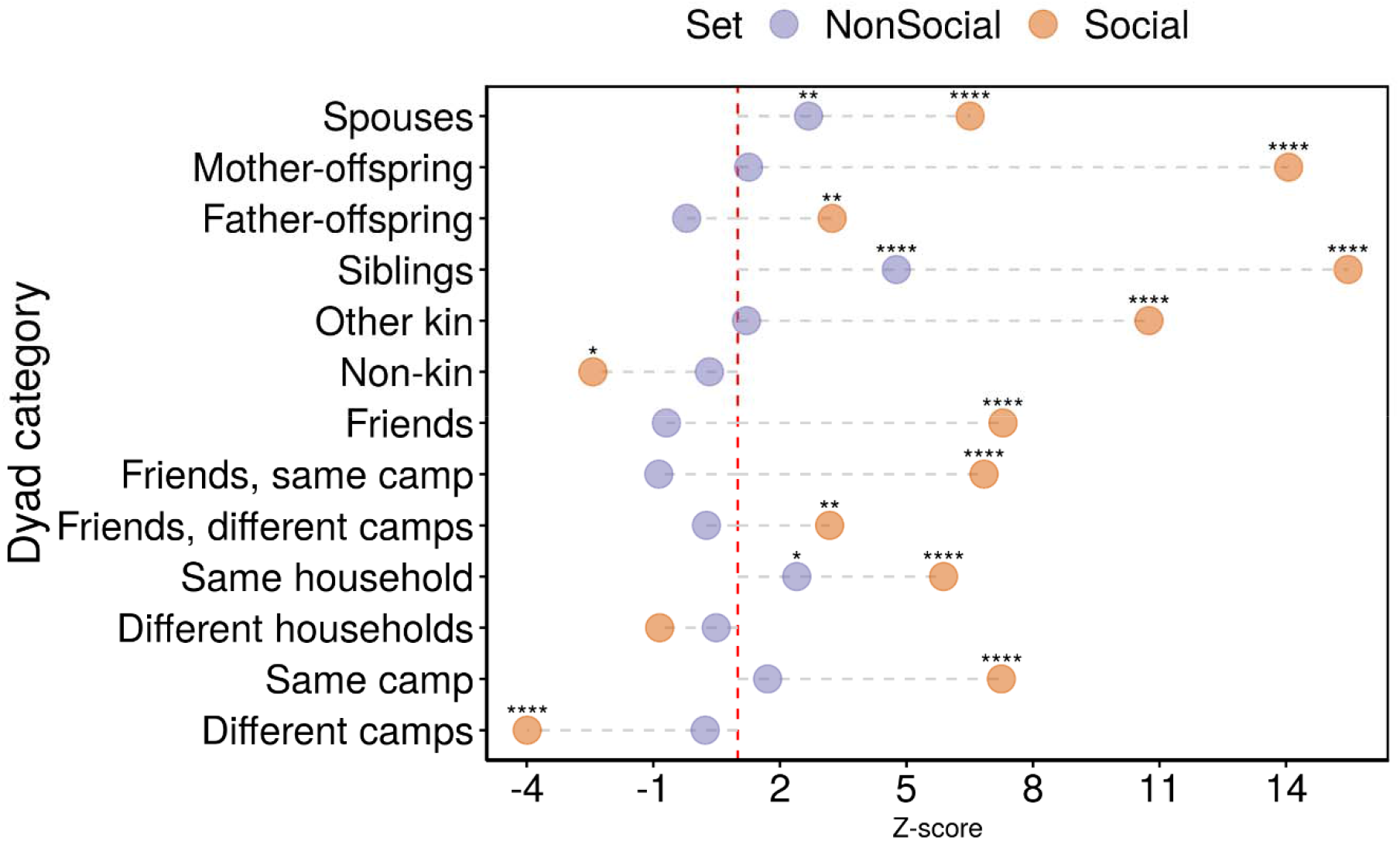
Effect of kinship, friendship and residence on dyadic bacterial sharing. Dyads were classified into kinship levels; same or different households; same or different camps; and between friends in the same or different camps. Dots show the z-score, or the standardised ratio of the mean link weight in real to randomised networks, in either social (orange) or non-socially transmitted bacteria (purple). Vertical red dashed line indicates a ratio of 1, or no difference between the number of shared bacteria in real and randomised networks. For socially transmitted bacteria, kinship, friendship and residence in the same household or camp are associated with significantly higher bacterial sharing than predicted from randomised networks of the same size and structure. By contrast, dyads from different camps or non-kin share significantly fewer bacteria than expected by chance; bacterial sharing in dyads from different households do not differ from randomised networks. For non-socially transmitted bacteria, the only dyadic categories significantly increasing bacterial sharing were siblings, spouses, and dyads from the same household (all of which share the same close environment). See Supplementary Tables 2 and 3 for values on mean weights for real and randomised networks. (****: FDR-adjusted P<0.0001; ***: P<0.001; **: P<0.01).

Results showed that some dyadic categories share significantly more socially transmitted bacteria than expected by chance. First, we observed higher bacterial sharing within the same household and camp than between different households and camps, an expected consequence of the Agta multilevel social structure. Kinship effects were also clear, with the highest levels of social microbiome sharing found in mother-offspring pairs, followed by siblings known interact every day in households and playgroups. High sharing between spouses also confirmed the importance of human pair bonding in microbial transmission. In addition, strong friendship links were also associated with increased bacterial sharing. Social bacterial sharing between friends in the same camp is as high as between close kin or within households. Friends in different camps also share a higher proportion of social bacteria than expected by chance, which is possibly a consequence of high between-camp mobility. By contrast, non-kin or individuals from different camps share fewer social bacteria than expected, further demonstrating the role of Agta friendships in the transmission of social bacteria across households and whole camps.

The same analysis performed instead on non-socially transmitted ASVs did not reveal significant effects on bacterial sharing from most dyadic categories, except for three types: spouses, siblings, and same household. A possible explanation is that some non-socially transmitted bacteria may be shared due to a common environment and diet in the same household. For example, we have shown in our parallel study^19^ that the proportion of meat versus rice in individual diets affects the composition of the oral microbiome. Therefore, similar diets may explain the presence of the same ASVs within the individuals of a household irrespective of social interaction levels. However, the effects of sociality and shared diets seem to be independent. This is shown by the fact that socially transmitted bacteria are equally likely to be related or not to diet: 13 socially transmitted genera were found also associated to diet (41 genera), whereas 23 were not (57 genera) (proportion test: chi-squared=0.44, P=0.51; See companion paper^19^ for further data and analysis). There we also show that host genotype correlates with the presence of certain ASVs19. While high genetic relatedness may play a role in bacteria sharing within households, none of the ASVs associated with the host genotype were present in the social microbiome. Therefore, our analyses seem to distinguish between the effects of social contact from shared environment or genes within dyadic types. Overall, the results show the roles of mobility and the multiple interaction channels created by multilevel sociality in social microbiome sharing, similarly to what is also observed in cultural transmission^6,18,26,34^.

### Frequency of social contact predicts social microbiome sharing

Although previous studies have investigated patterns of bacterial sharing in human groups, they have often been unable to comprehensively characterize transmission patterns due to limited information on social contact^35^. In order to obtain a full picture of individual contact and exposure levels, we built social networks based on proximity data from four camps and two multi-camp locations (Figure 3a-b). Overall, Agta social networks reveal a multilevel structure of households (mostly consisting of strong kin links) connected by a few strong links (mostly among unrelated friends) in each camp, and in the case of multi-camp groups, camps interconnected mostly due to visits among friends. We also observed equality of interactions within and between sexes and across age groups^6,18^. This pattern creates multiple channels for social transmission of bacteria both within and between camps, between close kin and unrelated individuals, and finally across whole multi-camp structures.

**Figure 3.**
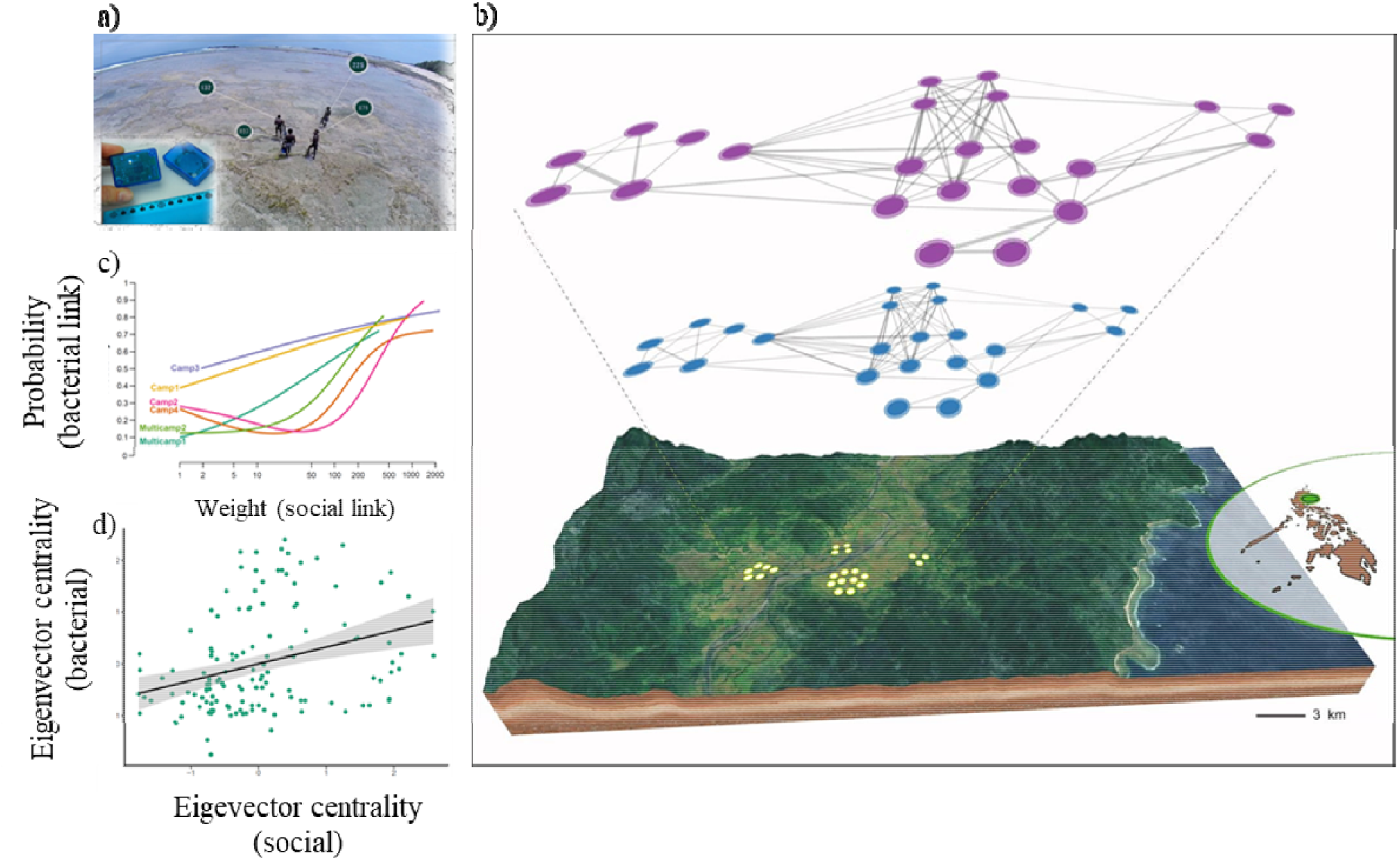
Characterisation of the social microbiome. a) Recording networks of social interactions using radio sensor technology. b) Reinforcement analysis estimates the probability of a link occurring in the bacterial sharing network (top layer, purple) based on the weight of the same link in the social contact network (bottom layer, blue). Network nodes (circles) represent the same Agta individuals in the bacterial sharing and social contact network. Panel displays networks from multi-camp 1 (23 individuals). Map shows geographical location of four camps interconnected by frequent migration. c) Probabilities of links in the Agta bacterial sharing network increase with their weights in the social contact network. Curves estimated by generalised additive modelling (binomial option). Data from four Agta camps and two multi-camps. d) Eigenvector centralities in bacterial sharing and social contact networks. Linear regression based on pooled data from four Agta camps and two multi-camp structures. Virtually similar results were obtained by including camp either as a fixed factor in a multiple regression (with or without interactions), or as a random factor (on intercept and slope) in a mixed effects linear regression.

To further assess whether increasing levels of social contact predict higher levels of sharing of socially transmitted bacteria, we applied reinforcement analysis^36^ (Figure 3b) to assess whether the social network predicts (or reinforces) the bacterial sharing network in each Agta camp. We calculated the conditional probability of each link between two individuals A and B in the bacterial network provided the same weighted link is present in the social network (Supplementary Table 4 and 5). For all four camps and two multi-camp structures, results showed that the weight of a dyadic link in the social network significantly predicts the probability of the same link occurring in the bacterial sharing network (Figure 3c and Supplementary Figure 2). Specifically, a larger dyadic weight in the social network implies a higher probability that the same individuals also share at least one socially transmitted ASV. For example, for multi-camp 1, while weak social network links (with weights under 10 recorded social interactions) show a probability below 20% of sharing any socially transmitted bacteria, strong links (over 200 recorded contacts) are associated with a probability above 70%. Overall, the results confirm that the Agta social microbiome is shaped by their social interactions.

### Hypersocial individuals are supersharers

We also investigated whether more socially interactive individuals exhibited higher social microbiome diversity. We calculated eigenvector centralities for all individuals in the bacterial sharing network, resulting in a significant and positive slope in a regression on eigenvector centralities from the same individuals in the social network (b=0.32, *P*=0.0001, *R*^2^ = 0.1, n=138, Figure 3d). We also identified 16 individuals ranked at the top quartile of eigenvector centralities in both networks as potential microbial ‘superspreaders’ or ‘superacquirers’, that is, “supersharers”. They do not stem from a specific age (7 to 68 years) or sex (six males, ten females), which is compatible with the egalitarian social structure of hunter-gatherers allowing individuals from any age or sex to be potentially central to social networks.

## Discussion

We have identified and characterized a socially transmitted fraction of the Agta hunter-gatherer oral microbiome. This fraction (7%) is surprisingly small, since in principle all 1980 identified ASVs could be orally transmitted between closely interacting people. Nonetheless, our results demonstrate a significant and independent role of social interactions on the transmission of oral microbiome, in addition to other factors such as shared environment (household and diet) and host characteristics (age, sex and genes) previously investigated in other hunter-gatherer populations^8–16^ and in the same Agta population^19^. The transmission of the 137 bacteria classified into the social microbiome seem to be facilitated by the extended sociality of hunter-gatherers and its various transmission channels, ranging from spouses to unrelated friends often residing in different camps. Together, reinforcement analysis, multiple channels of social interaction, and supersharers show that social microbiome sharing is strongly shaped by hunter-gatherer multilevel sociality. From an evolutionary perspective, sociality has considerably changed from our closest ape relatives to ancestral humans, when adopting a hunter-gathering lifestyle meant exhibited higher frequency of social contact with unrelated individuals, larger networks of extended kin across large geographical regions, and more egalitarian interactions between and within sexes and across ages. Such changes may have affected patterns of pathogen transmission and affected the human microbiome as observed in current hunter-gatherers. As with our study of the Agta, future research should collect data on both social networks and social microbiomes from the same populations of non-human apes; such a dataset would provide a comparative basis for analysing of the role of social evolution on the human social microbiome. Hunter-gatherer social networks are efficient systems of cultural transmission, and its specific channels organised around kinship, friendship and camp interconnectivity are central for the organisation of between-household cooperation, food sharing and social learning^24,26^. Agta mothers with higher social network centrality enjoy increased access to help and reproductive success^7^, but our results have shown that efficient networks may also facilitate the spread of infectious diseases, and hence significantly affect the structure and composition of the Agta microbiome. Crucially the frequency of pathogenic bacteria is much higher in the socially than in the non-socially transmitted fraction of the Agta oral microbiome. Together with the association between hypersocial individuals and increased bacterial sharing, this suggests a tradeoff between potential fitness benefits and costs of increased pathogen transmission. We conclude that the predominantly pathogenic oral social microbiome we identified in hunter-gatherers may be at least partially the outcome of a tradeoff between the advantages of multilevel sociality and the cost of infectious disease.

## Methods

### Ethnographic data collection

#### Agta demography

Ethnographic data collection took place over two seasons in April-June 2013 and February-October 2014. We censused 915 Agta individuals (54.7% male) across 20 camps. For the current study we selected four camps and two multi-camp structures where we collected data both on proximity networks and saliva samples. Accurate ages were estimated following relative aging protocols^37^. Relatedness (biological and affinal) was based on household genealogies. To resolve inconsistencies, we took either the genealogy from the most knowledgeable individual (i.e. mother over aunt) or the genealogy that reduced other inconsistencies (i.e. discarding six-month interbirth intervals). Genealogies contained 2953 living and dead Agta. We used the *R* packages *pedigree, kinship2*, and *igraph* to measure consanguineous relatedness (*r*)^34,38^. For comparative purposes, we obtained 14 saliva samples from neighbouring Palanan farmers, making sure individuals were unrelated by directly asking.

#### Bayaka demography

Ethnographic data collection took place over two seasons in April-June 2013 and February-October 2014. We collected saliva samples from 21 individuals for microbiome analyses.

#### Ethics

This study was approved by UCL Ethics Committee (UCL Ethics code 3086/003) and carried out with permission from local government and community members. Informed consent was obtained from all participants, after group and individual explanation of research objectives in the indigenous language. A small compensation (usually a thermal bottle or cooking utensils) was given to each participant. The National Commission for Indigenous Peoples (NCIP), advised us that the process of Free Prior Informed Consent with the tribal leaders, youth and elders would be necessary to validate our data collection under their supervision. This was done in2017 with the presence of all tribal leathers, elders and youth representatives at the NCIP regional office, with the mediation of the regional officer and the NCIP Attorney. The validation process was approved unanimously by the tribal leaders, and the NCIP, and validated the full 5 years of data collection.

### Oral microbiome analysis

#### Microbial DNA extraction and 16S rRNA gene sequencing

A total of 190 saliva samples were selected from Agta hunter-gatherers (n = 155) and Palanan farmers in the Philippines (n = 14), and Bayaka hunter-gatherers from the Congo (n = 21). Microbial DNA was extracted following the protocol for manual purification of DNA for Oragene·DNA/saliva samples. The 16S rRNA gene V3-V4 region was amplified by PCR with primers containing Illumina adapter overhang nucleotide sequences. All PCR products were validated through an agarose gel and purified with magnetic beads. Index PCR was then performed to create the final library also validated through an agarose gel. All samples were pooled together at equimolar proportions and the final pool was qPCR-quantified before MiSeq loading. Raw Illumina pair-end sequence data were demultiplexed and quality-filtered with *QIIME* 2 2019.1^39^ and *DADA2*^40^, which generates single nucleotide exact amplicon sequence variants (ASV or ESV). ASVs are biologically meaningful as they identify a specific sequence and allow for higher resolution than operational taxonomic units (OTUs)^41^ or clusters of sequences above a similarity threshold, and thus an ASV is equivalent to a 100% similar OTU. Taxonomic information was assigned to ASVs using a naïve Bayes taxonomy classifier against the *SILVA* database release 132 with a 99% identity sequence^42^.

Reads outside the kingdom Bacteria or assigned to mitochondria or chloroplasts were removed. Phylogenetic analyses aligned sequences with *MAFFT*^43^ and generated a rooted phylogenetic tree with *FastTree2^44^* using default settings via *QIIME* 2. We generated an Alpha rarefaction curve with R package *vegan* to confirm that sample richness had been fully observed (Supplementary Figure 3).

Samples with extremely low number of reads (8000) were removed. This resulted in 6409 ASVs (later reduced to 1980 ASVs present in at least two individuals and abundance of at least 10 counts per individual) and 173 individuals: 138 Agta, 21 Bayaka, and 14 Palanan farmers.

#### Identification of the Agta social microbiome

In our Agta sample, we first selected a set of strong social links (top 25% of the weight distribution from each camp and multi-camp). For each ASV, we calculated the proportion of strong links 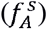 where a given ASV A was present. Next, we calculated the same proportion in the complementary set of 75% weak social links 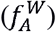. We then computed for each ASV A the score 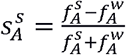, or normalised difference between the two proportions. This score can be paired with the z-score 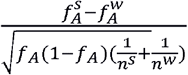 which quantifies the deviation from the null hypothesis that the two proportions are equal, with *n^s^* and *n^w^* as respectively the numbers of strong and weak links and *f_A_* as the proportion of total links that share ASV A. We then selected as affected by social interaction 137 ASVs with s >0.5 and P< 0.05. P-values were adjusted by False Discovery Rate (FDR).

We performed a sensitivity analysis to investigate the consequences of varying the threshold defining strong versus weak social links in the Agta social network. Instead of 137 ASVs resulting from selecting dyads at top 25% of the weight distribution, we obtained:

Top 45%: 166 ASVs
Top 35%: 156 ASVs
Top 25%: 137 ASVs
Top 15%: 138 ASVs
Top 5%: 99 ASVs

The list shows that whether the strong set consists of a very reduced number dyads with very strong weights (top 5%), or instead includes nearly half the dyads (top 45%), the number of ASVs significantly responsive to social contact varies from 99 (5%) to 166 (8%), representing a small fraction of the total of 1980 ASVs found in the Agta. Therefore, setting the threshold at the top 25% did not affect our results and conclusions.

### Agta social network data, construction and analysis

#### Mote devices

Motes are wireless sensing devices storing all between-device communications within a specified distance and have been described in detail elsewhere^6,7,18,45^. We used the UCMote Mini (with a TinyOS operating system) sealed into wristbands or belts, labelled with a unique number, and identified with coloured string to avoid accidental swaps. Motes require no grounded infrastructure and collect interactions even when individuals are away from camps. Individuals arriving at a camp after the start of data collection were given a mote and entry time was recorded, while those leaving a camp before the end of data collection had their exit time recorded. To prevent swaps individuals were checked twice daily, and mote numbers were checked upon return. Any swaps were later corrected by reassigning data to the correct individuals.

Data were later downloaded via a PC side application in *Java*. Data were limited to 5am-8pm. We ran raw data through a stringent data-processing system in *Python* to prevent data corruption. Data were matched to ID numbers and start-stop times of each mote. The result was a matrix with the number of recorded beacons for all possible dyads and their weights.

For the camp-level experiment, all individuals from four camps wore motes from five to seven days. Each device sent a message every two minutes that contained its unique ID, a time stamp and the signal strength. Messages are stored by any other mote within a three-meter radius, a frequently used threshold^46,47^. For the multi-camp experiment, adult individuals from two areas (consisting of seven and three camps respectively) wore motes for one month.

#### Effect of dyad category on bacterial sharing

The bacterial sharing network was constructed by defining link weights as the number of social bacteria shared by two individuals. Dyads in the network were classified into: i) levels of kinship (mother-offspring, father-offspring, siblings: *r*=0.5, other kin: *r*=0.25 or r=0.125, non-kin: *r*=0.0625 or lower, spouses, friends: defined as non-kin at the top 25% distribution of social dyadic weights, and other non-kin) and ii) residence (same or different camp, same or different household). Mean weights were calculated for each dyadic category (Supplementary Table 2). Then, we produced 1000 network randomisations based on a single-step ID swap between nodes. For example, if dyad 1 consisted of two spouses in the real network, randomisation preserved dyad 1 and its weight, but randomly replaced the two nodes (potentially changing the dyadic classification to siblings, friends, etc.). We calculated the mean weights for each dyadic category in the 1000 randomised networks, and then calculated one-sample t-tests with the mean weight in the real network as the test value. We repeated the analysis for non-social bacteria (Supplementary Table 3).

#### Reinforcement analysis

In a multilayer network, reinforcement analysis measures the overlap in links between different layers to quantify the probability of finding a link on a layer conditioned on the weight of the same link on another^48^. Reinforcement between two network layers *α* and *α*’, or *P*(*α*’|*α*), is defined as

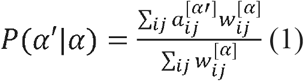

where 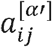 is the adjacency matrix of conditioned layer *α*’, and 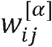 is the adjacency matrix of the conditioning layer *α*. We split the weight of social links into three tertiles, and computed equation (1) for each. We obtained increasing values of reinforcement from the lower to the higher tertile, providing evidence of an effect of social contact on bacterial sharing.

### ASV classification

#### ASV diversity metrics

To distinguish between the effects of lifestyle and shared ecology on the microbiome, we compared the diversity of the oral microbiome of Agta hunter-gatherers with neighbouring Palanan farmers in the Philippines and with Bayaka hunter-gatherers in Congo. Using the 6409 ASVs dataset we calculated the number of observed ASVs in each population with *R* package *Phyloseq* (version 1.30.0) and the Faith’s Phylogenetic Diversity index was calculated with *R* package *picante* using the generated rooted phylogenetic tree. To estimate the number of shared ASVs in the Agta, Bayaka and Palanan farmers, we sampled a random subset of 10 samples for each population without replacement, calculated the shared ASVs between the populations and repeated this procedure 100 times. Global differences between groups and pairwise comparisons were assessed by Kruskal-Wallis and Wilcoxon Rank Sum tests respectively and plotted by the *R* package *ggpubr*. Pairwise p-values were adjusted by False Discovery Rate (FDR).

#### Classification of oral bacteria as pathogens

ASVs were classified as oral pathogens if they have been reported as etiological agents of periodontitis^49,50^ or dental caries^51,52^. For gum disease pathogens, these included the classical “red” and “orange” complex of periodontal pathogens and the recent update by Pérez-Chaparro and cols^50^ based on systematic review and metaanalysis. For caries pathogens, the list of active microorganisms detected through metatranscriptomics of cavities was used. Common oral commensals potentially causing endocarditis or systemic infections in immunocompromised patients only were not considered pathogens. Bacteria reported as etiological agents of lower respiratory infections (e.g. pneumonia, whooping cough, bronchitis, or sinusitis) and biofilm-mediated infections (e.g. lactational mastitis, medical implant biofilm infections, chronic lung infections, osteomyelitis or chronic wounds) were also considered pathogens, including organisms present in healthy carriers^53,54^. Bacteria causing urinary tract infections or sexually transmitted diseases transiently found in the oral cavity were also considered pathogens^55^. Bacteria were classified as “oral” if detected in more than 10% of the population in oral samples according to the Human Oral Microbiome database. If a bacterial species or genus had been isolated from the oral cavity of an animal, it was also classified as oral.

For assignment of bacteria to pathogenic or non-pathogenic, we used species-level ASVs, given that there are multiple cases where different species from the same genus had a different assignment. If taxonomic classification of the ASV was only possible at the genus level, it was considered a pathogen if: i) >90% of named species within the genus were pathogenic, or ii) the genus included a major pathogenic species but the remaining species within the genus were not classified as oral by the Human Oral Microbiome Database^56^. ASV with a top hit to a sequence classified as “Oral taxa” in databases but without a species assignment were not considered named species and were discarded from the analysis. Cases where taxonomic classification of the ASV was only possible at the family level or higher were also discarded.

## Acknowledgements

This project was funded by Leverhulme Trust Grant RP2011-R 045 to A.B.M. J.B. received grant PID2019-110933GB −I00/AEI/10.13039/501100011033 by the Agencia Estatal de Investigación (AEI), Ministerio de Ciencia, Innovación y Universidades (MCIU, Spain) and support from Secretaria d’Universitats i Recerca del Departament d’Economia i Coneixement de la Generalitat de Catalunya (GRC 2017 SGR 702). Part of the “Unidad de Excelencia María de Maeztu”, funded by the AEI (CEX2018-000792-M), that granted the microbiome analysis by the Servei de Genòmica, Universitat Pompeu Fabra. A.M is funded by grant RTI2018-102032-B-100 from Ministerio de Ciencia, Innovación y Universidades (MCIU, Spain).

## Data availability

16S amplicon data (EGAS00001005317) are deposited at the European Genome-phenome Archive (EGA), which is hosted at the EBI and the CRG. Data at the individual level on age, kinship relationships, household composition, camp assignation and social contacts that support the findings of this study are available on request from the corresponding author (A.B.M.). The individual data are not publicly available due to information that could compromise research participant privacy.

## Code availability

Source code and data for visualization are available at https://doi.org/10.5281/zenodo.6338840

