## Supplementary Figures for "Hunter-gatherer oral microbiomes are shaped by contact network structure"

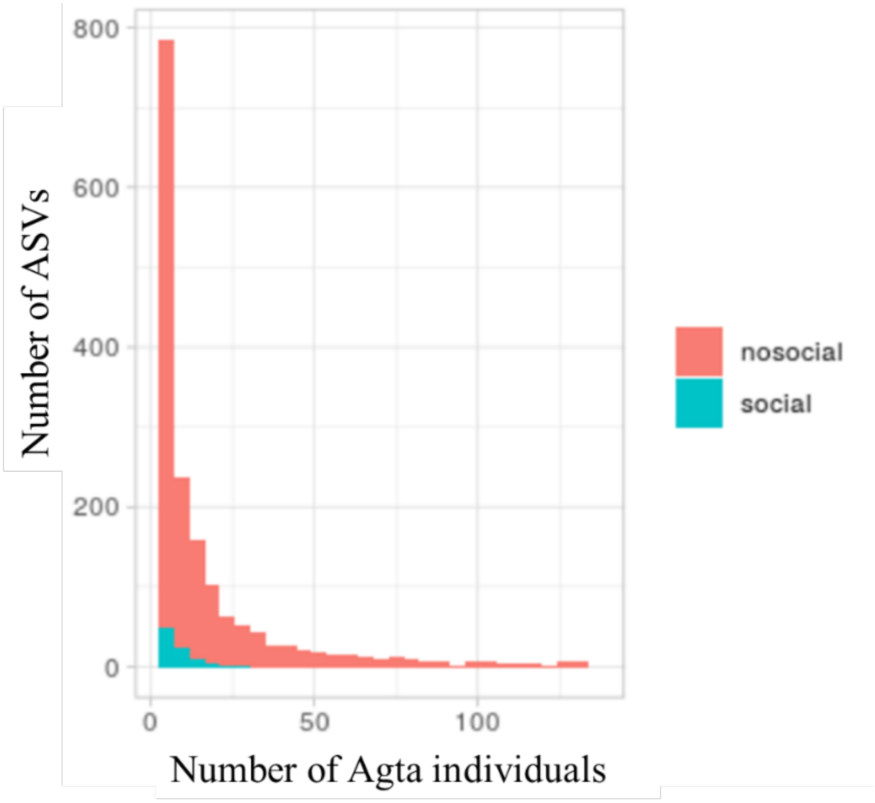


**Supplementary Figure 1. Number of socially and non-socially transmitted ASVs per Agta individual.** Plot shows that around 50 (out of 137) socially transmitted ASVs were found in only one Agta individual (out of 138); very few were found in more than 25 individuals. For non-socially transmitted ASVs, about 800 (out of 1843) were found in only one individual; overall, only 36 bacteria were found in more than 100 individuals.

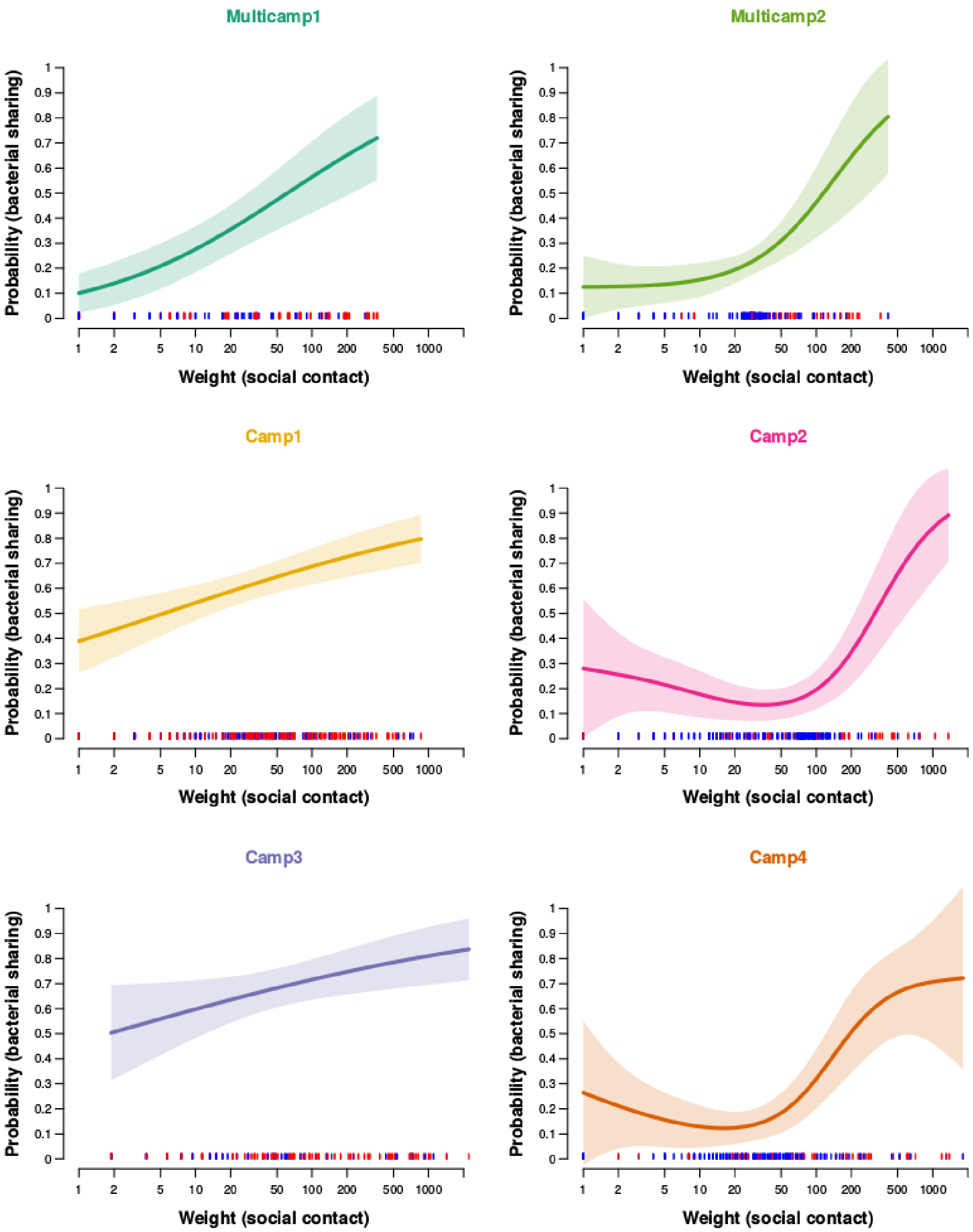


**Supplementary Figure 2. Reinforcement analysis.** Probabilities of links in the Agta bacterial sharing network in relation to their weights in the social contact network. Curves estimated by generalised additive modelling (binomial option). Data from two multi-camps and four Agta camps. Lines indicate single data points.

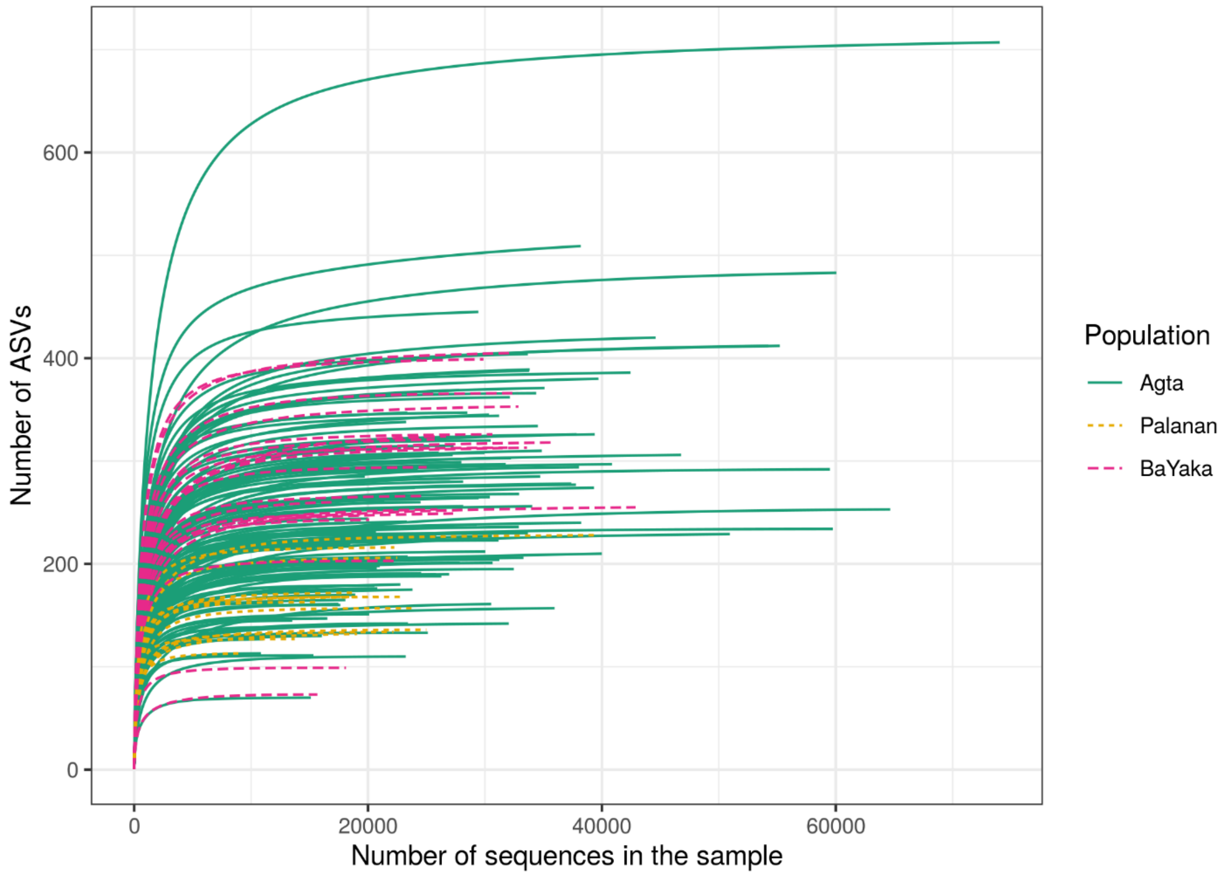


**Supplementary Figure 3. Rarefaction curves calculated at an interval step of 50 for each sample.** Each line shows the cumulative number of different ASVs found in a sample based on the number of sequences sampled.
